# Plant Face: Machine learning decodes genetic, environmental and developmental imprints in leaf appearance

**DOI:** 10.64898/2025.12.01.691347

**Authors:** Jie Li, Yongxian Chen, Zenghong Gao, Xuan Zhou, Zhan Xu, Yin Zhang, Kaichi Huang, Guangwei Ma, Yong-Juan Zhao, Jian Li, Xin Liu, Yabin Guo

## Abstract

Leaf appearance is a crucial plant phenotype. However, traditional methods for extracting this information are inefficient, limiting its full utilization. Deep learning based on convolutional neural networks (CNNs) enables us to capture previously inaccessible information from images. In this study, we made the surprising discovery that the leaf appearance of each individual plant is unique. Using deep learning, leaves from one plant could be efficiently distinguished from those of another plant of the same species and cultivar. We term this phenomenon the “*Plant Face*” and suggest the potential to develop a “plant face recognition system,” analogous to human facial recognition. We also applied similar methods to study the relationship between leaflet appearance and their position on compound leaves, leaf bilateral symmetry, and differences in leaves from twining stems with different chirality. These results collectively indicate that plant genetic characteristics, growth conditions, and developmental features can be stored within their appearance. With appropriate decoding, leaf appearance is poised to play an increasingly important role in phenomics.

**Significance:** The saying “no two leaves in the world are identical” holds philosophical significance, as such variation encompasses considerable contingency and randomness. Here, we assert that **no two trees have identical leaves**; meaning that even for plants of the same species and cultivar, the leaf morphology of each individual plant is distinct at the population level, even though single leaves may overlap in appearance.

Genetic, environmental, and developmental information is recorded in some manner within the phenotypic appearance of leaves. With advancements in computational technologies like artificial intelligence, this information can now be decoded.

**Highlights:**

- The leaves of each individual plant are statistically unique.
- The relationship between leaflet appearances in compound leaves hints at their developmental patterns.
- Leaves are not necessarily bilaterally symmetric in a statistical sense.
- Leaves from stems with different chirality (twining direction) exhibit distinct appearances.

## Introduction

Plant leaf appearance is undoubtedly a crucial phenotypic trait. Traditional morphometric methods, however, often rely on limited manual feature extraction (e.g., leaf length, width, area, aspect ratio, color), which offer limited information density and struggles to capture high-dimensional, nonlinear, and subtle phenotypic patterns [1, 2]. This significantly restricts the practical application of leaves as a key component of the phenome. Recent advances in artificial intelligence, particularly deep learning, provide a revolutionary solution to this challenge [3]. Models like convolutional neural networks (CNNs) can automatically learn hierarchical feature representations from raw images, eliminating the need for manual feature design based on prior knowledge [4]. This “end-to-end” learning paradigm demonstrates remarkable potential for discovering complex patterns imperceptible to the human eye or traditional algorithms, as evidenced by its remarkable success in medical image analysis and face recognition [5, 6].

Today, computer-based recognition of human faces is commonplace, and similar technology has been applied to animals, distinguishing individuals based on facial features [7, 8], fur patterns, spots, etc. [9, 10]. In plant science, deep learning has been widely used for species classification [11, 12] and disease detection [13–15]. However, there are no reports in plants analogous to human or animal individual identification via comparison of photographs of a specific organ. This might be due to the default assumption that organ appearance differences between plant individuals are less distinct than those in animals. In reality, the possibility should not be overlooked that each plant individual might also possess uniquely appearing organs, which are relatively stable, observable, and identifiable. Accordingly, we hypothesize that even for plants of the same species and cultivar, their genomes are not completely identical; coupled with potentially varying growth environments, their gene expression profiles also differ. These characteristics are recorded in some manner during leaf growth into the leaf’s appearance. That is, each plant has a unique leaf appearance, akin to a human face, which we term the “Plant Face.” If this hypothesis holds, deciphering these subtle, “beyond human visual resolution” faces could open new avenues for understanding plant individuality, developmental plasticity, and micro-mechanisms of environmental adaptation.

In this study, we used deep learning to compare leaf appearances from multiple individual plants of various species, providing preliminary confirmation of the hypothesis: leaves from different individual plants of the same species are distinguishable. Furthermore, we used similar methods to investigate differences in leaflet appearance at different positions on compound leaves, leaf bilateral symmetry, and differences in leaves from twining stems of different chirality in the vine *Mikania micrantha*, all yielding satisfactory results.

## Results

### Identification of Leaves from Two Trees

To test our hypothesis, we began by comparing leaves from two trees. We selected a common ornamental tree in Southern China, the Cockspur Coral Tree (*Erythrina crista-galli*), as our subject. Its leaves are trifoliate. As shown in Fig. 1A, we only used the central leaflet (position C) for comparison. Approximately 200 leaves were collected from each of two trees (#0 and #1). After photographing, 60% of the images were randomly selected for training, with 20% each for validation and testing. Surprisingly, after 57 epochs of training (Fig. 1B&C), both validation and test accuracy for distinguishing leaves from the two trees reached 100% (ROC AUC=1, P= 8.67E-22) (Fig. 1D-F). Dimensionality reduction visualizations (t-SNE and UMAP) of image similarity distributions also showed successful separation of the two image groups (Fig. 1G&H). The average images computed by overlaying all photos from each group also revealed visible differences (Fig. 1I). We performed other pairwise comparisons between *E. crista-galli* individuals. Supplementary Fig. 1 shows results for #0 vs #2 and #1 vs #2, where validation and test accuracy also reached or approached 100%.

**Fig. 1.**
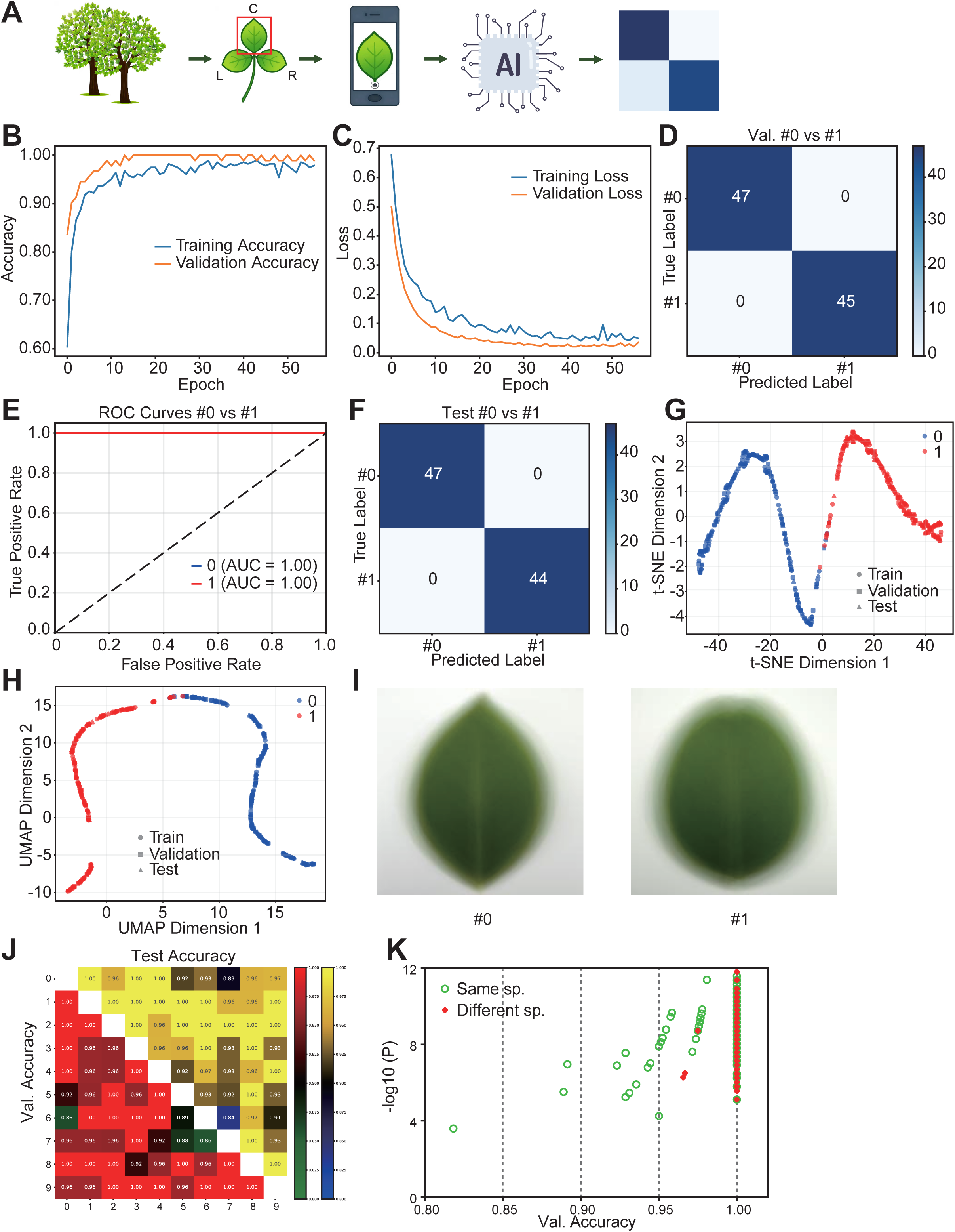
Comparison of leaves from two plants of *Erythrina crista-galli*. A, Study design: collecting the terminal leaflet (position C) from the middle of the compound leaves, photographing them, and training a learning model using convolutional neural network (CNN). B, Training history: changes in Accuracy (B) and Loss (C) relative to Epoch. D, Confusion matrix of validation. E, ROC curve of the model. F, Confusion matrix of test. G, t-SNE dimensionality reduction visualization of image similarity distributions. H, UMAP dimensionality reduction visualization of image similarity distributions. I, Average images of leaves from the two plants. J, Results of pairwise comparisons of leaves from 10 plants of *E. crista-galli*; the heatmap shows the Validation Accuracy in the lower left triangle and the Test Accuracy in the upper right triangle. K, Scatter plot showing the Validation Accuracy and statistical significance (-log10(P)) for each of the 276 pairwise comparisons performed on leaves from 24 individual plants belonging to 3 different species, 14 of *E. crista-galli,* six of *Triadica sebifera* and four of *Osmanthus fragrans*.

Subsequently, we conducted batch validation. We performed pairwise comparisons for leaves from 10 *E. crista-galli* individuals. Considering time constraints, we used only 60-70 images per pair (Fig. 1J). The heatmap’s lower left triangle shows validation accuracy, with over half (24/45) reaching 100%; the upper right triangle shows test accuracy, with 21/45 at 100%, and the vast majority >90%. We then conducted a larger-scale test involving 14 *E. crista-galli*, 6 Chinese Tallow Trees (*Triadica sebifera*), and 4 Sweet Osmanthus (*Osmanthus fragrans*) individuals, totaling 24 plants and 276 pairwise combinations. Results showed that for within-species comparisons, 79/112 validation accuracies were 100%; while for between-species comparisons, only 3/116 were not 100% (Fig. 1K).

Although such results were expected by our hypothesis, we were nonetheless surprised by the high degree of distinguishability. This strongly suggests that each tree’s leaves are, at least statistically, unique. Even leaves from two plants of the same species and cultivar are highly distinguishable, or colloquially, “no two trees have identical leaves.”

### Identification of Leaves from Multiple Trees

Building on distinguishing leaves from two trees, we further attempted to distinguish leaves from multiple trees. We used the central leaflet (C) from 10 *E. crista-galli* individuals and trained a model using a similar approach. Results showed successful model training (Supplementary Fig. 2A), with validation and test accuracy both exceeding or approaching 80% (Fig. 2A&B, P=0, Supplementary Fig. 2C&D). The minimum ROC AUC was 0.97 (Fig. 2C). t-SNE and UMAP clustering of image similarity distributions also showed successful separation among the photo groups (Fig. 2D&E). Fig. 2F displays the average image for each group, helping us understand the systematic differences between leaves of different plants, which might be more significant than commonly perceived. This also indicates that machine learning allows us to detect previously unnoticeable or at least unquantifiable phenotypic differences.

**Fig. 2.**
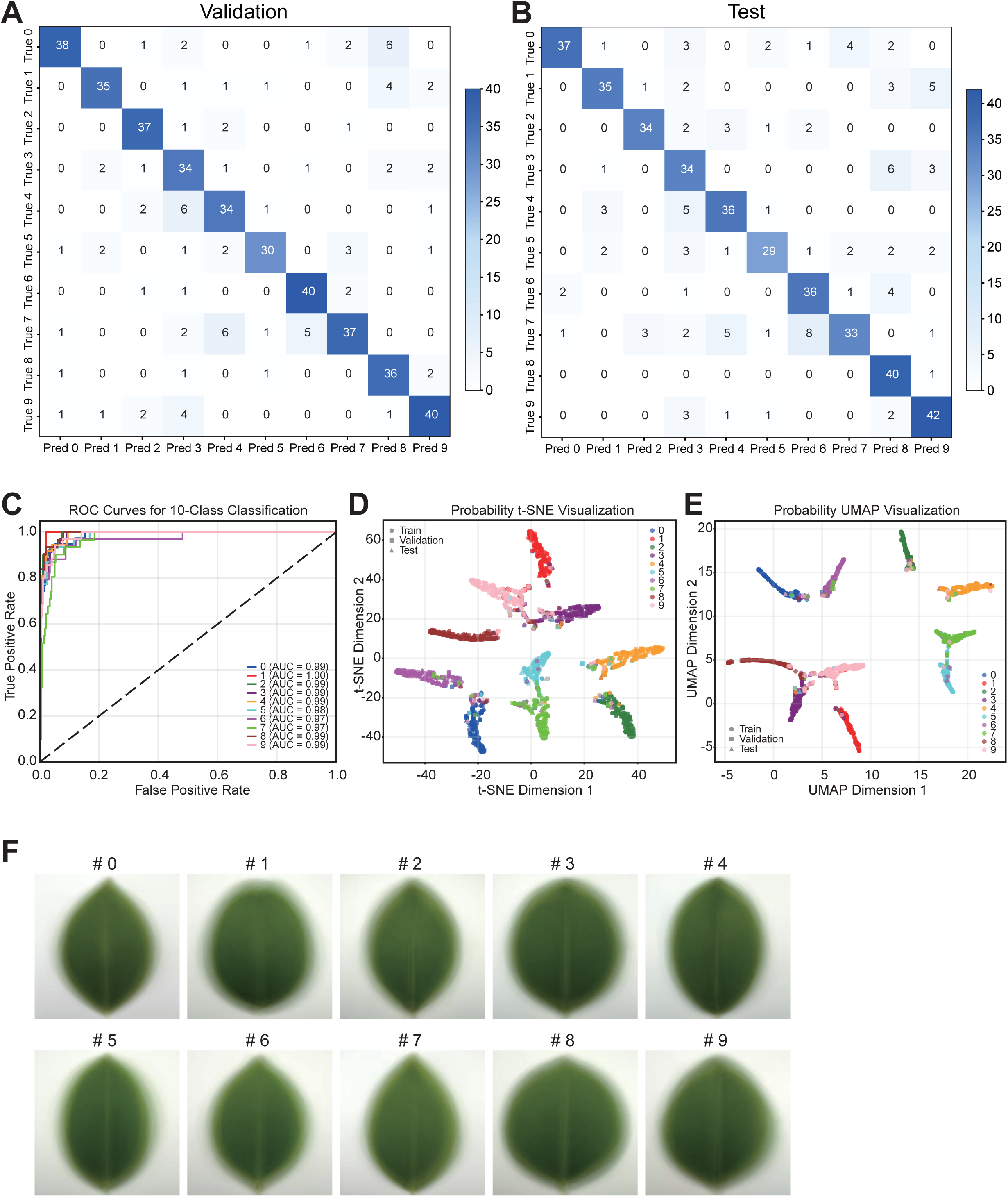
Collective comparison of leaves from 10 E. crista-galli plants. A, Confusion matrix of Validation Accuracy (80.1%). B, Confusion matrix of test Accuracy (77.9%). C, ROC curve of the model. D, t-SNE dimensionality reduction visualization of image similarity distributions. E, UMAP dimensionality reduction visualization of image similarity distributions. F, Average images of leaves from the ten plants.

Naturally, ten-class identification is far more challenging than two-class. If the number of plants increases further, the identification difficulty will also increase, but this is a technical challenge. The difficulty of multi-plant identification does not negate the objective existence of inter-plant differences or the fact that each tree’s leaves are unique.

### A Plant Face Recognition Machine

The previous results suggest that plant leaves, in a sense, are like human faces, enabling distinction between individuals. Therefore, we use the evocative term “plant faces” to describe the leaves. However, a person has only one face, while a plant has many leaves, which are not identical. Thus, “plant face” refers not to a single leaf, but to the systematic or statistical characteristics of a plant’s leaves. Given that the single-leaf accuracy in the ten-plant comparison reached ∼80%, this implies that if we identify five or more leaves at once, the accuracy could approach 100%. Based on this premise, we developed a “*Plant Face Recognition Machine*”.

Unlike previous comparisons, the recognition machine employs an interactive, dynamic approach. Users can enroll and test incrementally and decide the number of test images per “face” (leaves per face). Enrolled images are saved as a database, and the program loads previously saved databases upon each run (Fig. 3A). As differences between leaves of different species are typically much greater than those between leaves of different plants within the same species, this recognition machine can also enroll multiple plant species without interference. We conducted batch tests using images from 50 individual plants belonging to 18 species (Fig. 3B, Supplementary Table 1), running three tests per scenario and ensuring enrolled photos were not used for testing. Fig. 3C shows that even with only one enrolled image and using 1 leaf per face for testing, accuracy exceeded 50%, far greater than the random guess rate of 2%. When enrollment reached 8 images, testing with 8 leaves/face yielded satisfactory results. With 32 enrolled images and testing with 8 leaves/face, accuracy reached 100%. Fig. 3D shows the similarity distribution from one test instance. Fig. 3E shows t-SNE clustering of similarities, where each plant species (same color) occupies distinct, non-overlapping regions, and each individual plant (dashed triangles) is also separated.

**Fig. 3.**
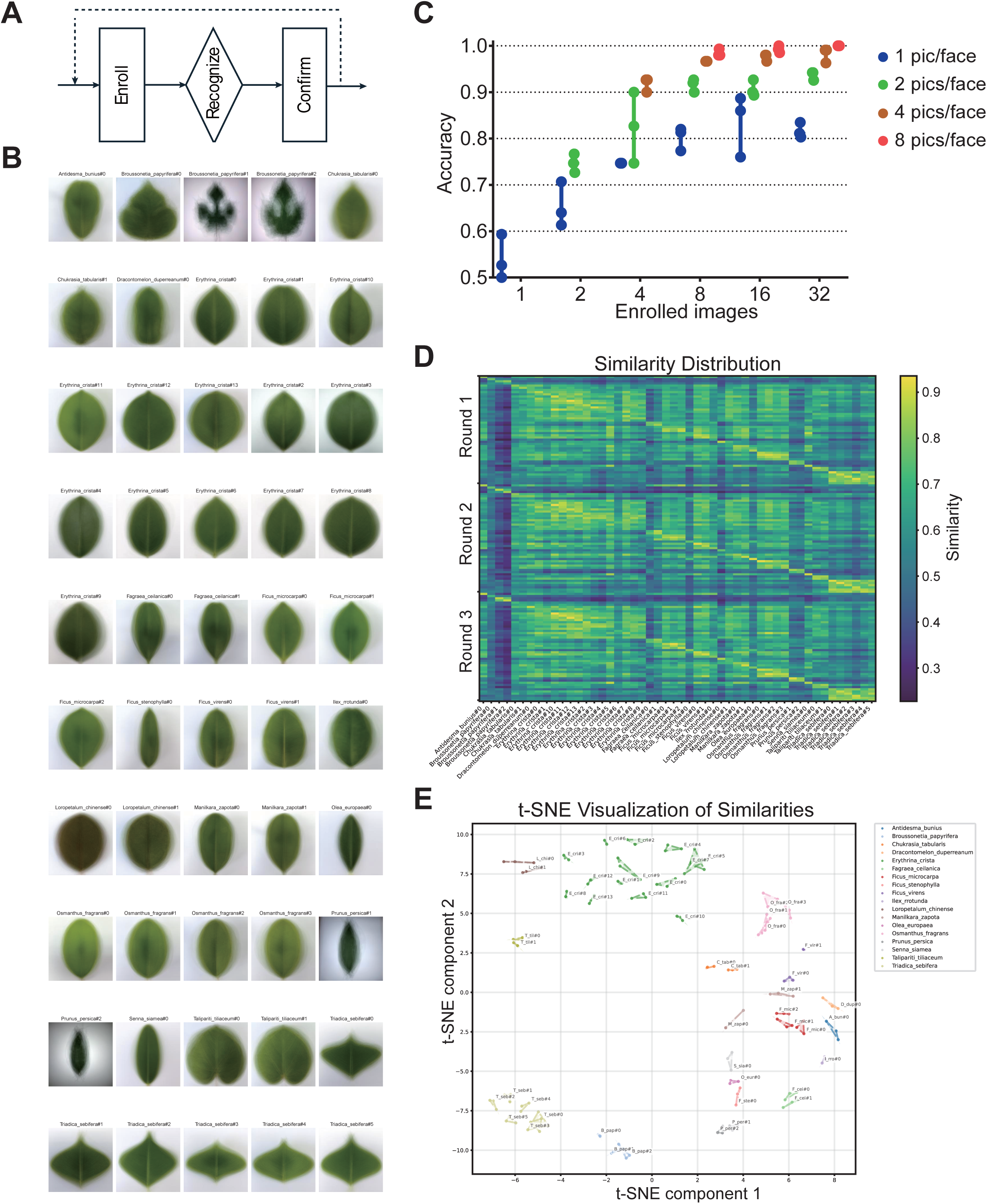
Plant “Face Recognition Machine”. A, Schematic diagram of the system workflow. B, Average leaf images from 50 individual plants belonging to 18 species. C, Batch test accuracy of the system, showing the accuracy achieved with different numbers of enrolled images per plant (x-axis) and different numbers of test images “pics/face” (represented by differently colored dots). D, Heatmap showing the accuracy distribution from one test scenario using 8 enrolled images and 8 test images per “face”. E, t-SNE dimensionality reduction visualization of the distribution from the previous panel; each color represents a different plant species, and dashed lines connect points representing the individual test of the same individual plant.

These results demonstrate that plants can be distinguished using a method analogous to human face recognition. The current version of this recognition machine is not yet as accurate as commonly used human face recognition systems, potentially due to: a) The maturity of human face recognition technology after decades of development; b) The dynamic nature of plant leaves, which continuously grow and abscise, unlike the relatively stable human face; c) The clonal origin of many horticultural plants, analogous to distinguishing identical twins.

### Relationship between Leaflet Appearance and Position on Compound Leaves

*E. crista-galli* leaves are trifoliate, each with three leaflets. Our initial study used only the central leaflet, but we were also curious whether the three leaflets exhibited systematic differences in appearance. Although the genomes of different leaflets are identical, during compound leaf differentiation, leaflets at different positions might experience different developmental environments, leading to differential gene expression and consequently different appearances. Therefore, we collected compound leaves from 6 *E. crista-galli* individuals, grouping leaflet photos by position into L (left), C (central), and R (right) (Fig. 4A&B). Each group was randomly split into train, validation, and test sets for model training (Supplementary Fig. 3A-E, Fig. 4C-E, Val. Accuracy=66.3%, P= 4.25E-12). L and R leaflets could be well distinguished from the C leaflet, but the left and right leaflets were often confused. This indicates highly similar developmental environments for symmetrically positioned leaflets, while the lateral leaflets experience significantly different developmental environments compared to the central leaflet.

**Fig. 4.**
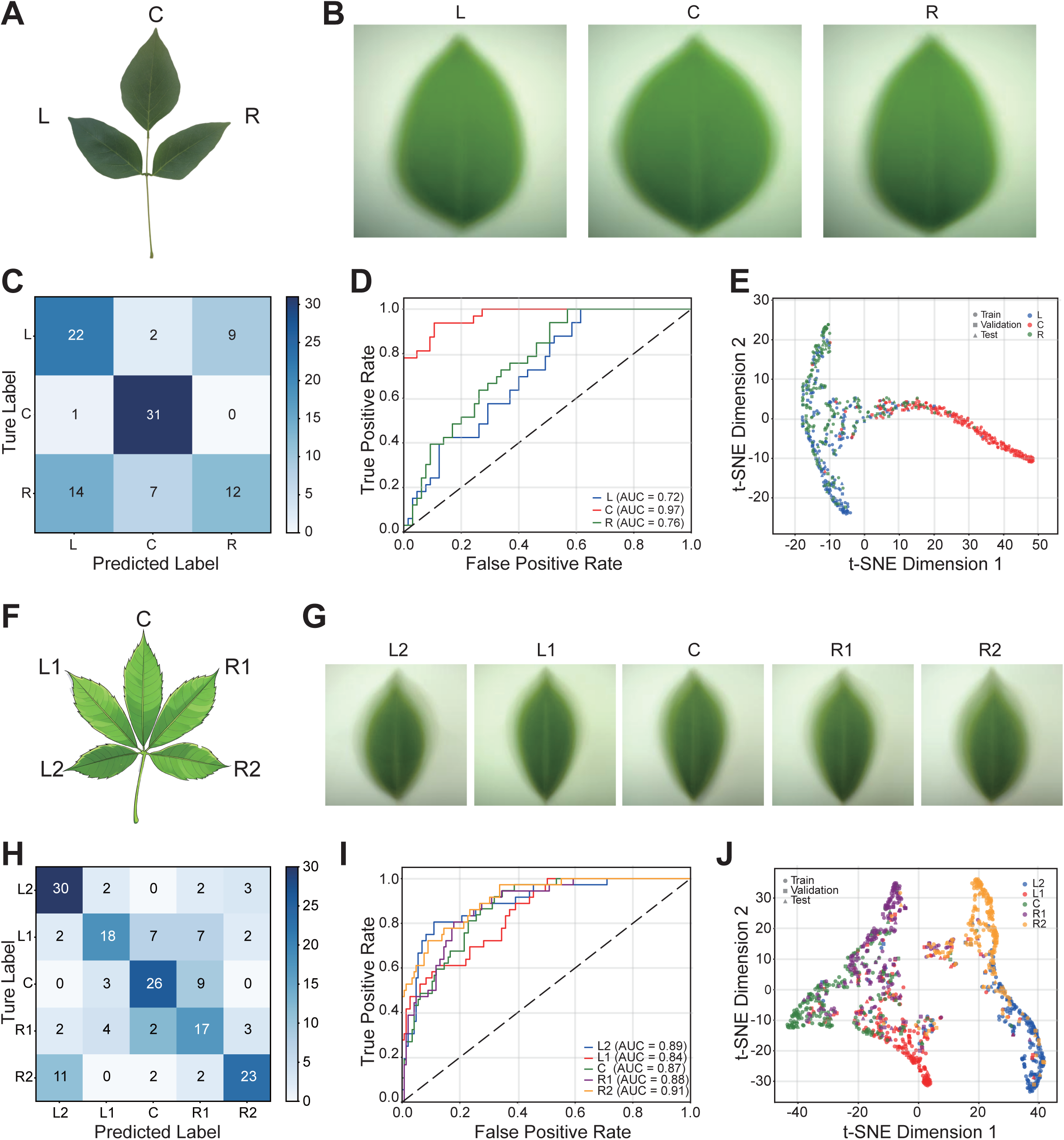
Relationships between the appearance of leaflets within compound leaves. A, A compound leaf of *E. crista-galli*. B, Average images of the three leaflets from different positions on the *E. crista-galli* compound leaf. C, Confusion matrix of validation Accuracy for CNN-based identification of leaflets from the three positions. D, ROC curve of the model. E, t-SNE dimensionality reduction visualization of image similarity distributions. F, A palmately compound leaf with five leaflets from *Ceiba speciosa*. G, Average images of the five leaflets from different positions on the *C. speciosa* compound leaves. H, Confusion matrix of Validation Accuracy for CNN-based identification of leaflets from the five positions. I, ROC curve of the model. J, t-SNE dimensionality reduction visualization of image similarity distributions.

Next, we considered more complex compound leaves. The palmately compound leaves of the Silk Floss Tree (*Ceiba speciosa*) typically bear 5-7 leaflets; here we used leaves with five leaflets (Fig. 4F&G) and studied them using methods similar to those mentioned before.

Identification results were slightly better than for the trifoliate *E. crista-galli* (Supplementary Fig. 3F-J, Fig. 4H-J, Val. Accuracy=64.4%, P= 2.29E-49). In summary: a) Adjacent leaflets were easily confused, especially L1 and R1 positions often misclassified as C; non-adjacent leaflets like L2 vs C and R2 vs C were rarely confused. b) Symmetrically positioned leaflets L2 vs R2 were also easily confused. This suggests the existence of a continuous gradient distribution (e.g., of hormones) during compound leaf development, making the developmental environments of adjacent leaflets more similar. Furthermore, this distribution is bilaterally symmetric, resulting in similar appearances for leaflets at symmetric positions.

### Leaf Bilateral Symmetry

Given that leaflets at symmetric positions on compound leaves were often confused, we specifically compared these leaflets. Results showed systematic differences between L and R leaflets in *E. crista-galli* (Fig. 5A&B, Val Accuracy=75.8%, P=2.85E-05, Supplement Fig. 4A). Since horizontal flipping was disabled during training and inference, any two images that are mirror reflections of each other—but not individually bilaterally symmetric—were treated as distinct by the model. Therefore, we mirrored the R leaflets (R’) and compared them with L, finding they became indistinguishable (Fig. 5C, Val Accuracy=53.0%, P=0.62, Supplement Fig. 4B&D). This indicates that the lateral leaflets are mirror images of each other. We compared the C leaflet with its mirror image (C’) and found them indistinguishable, indicating it is bilaterally symmetric itself (Fig. 5D, P=1, Supplement Fig. 4C&E). These results demonstrate that the entire *E. crista-galli* compound leaf is bilaterally symmetric.

**Fig. 5.**
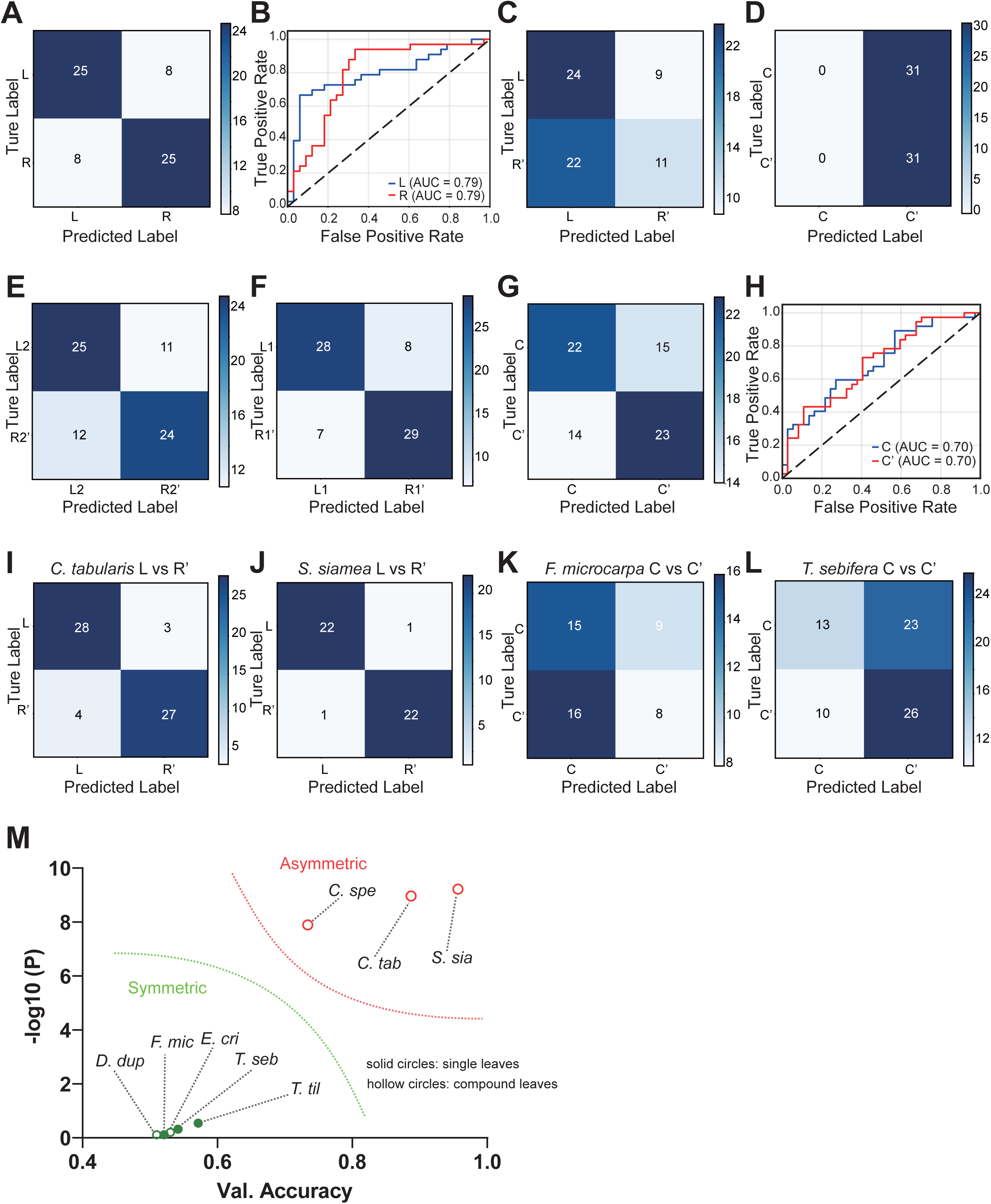
Leaf Symmetry. A, Confusion matrix for distinguishing between L and R leaflets in *E. crista-galli*. B, ROC curve for distinguishing *E. crista-galli* L and R leaflets. C, Confusion matrix for distinguishing between L and the mirrored images of R (R’) in *E. crista-galli*. D, Confusion matrix for distinguishing between the C leaflets and its mirror images (C’) in *E. crista-galli*. E, Confusion matrix for distinguishing between L2 and the mirrored R2’ in *C. speciosa*. F, Confusion matrix for distinguishing between L1 and the mirrored R1’ in *C. speciosa*. G, Confusion matrix for distinguishing between the C leaflets and their mirror images (C’) in *C. speciosa*. H, ROC curve for distinguishing *C. speciosa* C and C’ leaflets. I, Confusion matrix for distinguishing between the mirrored left leaflets (L’) and the right leaflets (R) in *Chukrasia tabularis*. J, Confusion matrix for distinguishing between the left leaflets (L) and the mirrored right leaflets (R’) in *Senna siamea*. K & L, Confusion matrices for leaf vs. mirrored leaf classification: *Ficus macrocarpa* (K), *Triadica sebifera* (L). M, Accuracy and statistical significance (-log10(P)) for distinguishing between the left and right halves of leaves from several plant species, used to assess whether the leaves exhibit bilateral symmetry.

We similarly examined the compound leaves of *C. speciosa*. L2 vs R2 (Fig. 5E, Accuracy=68.1%, P=0.01) and L1 vs R1 (Fig. 5F, Accuracy=79.2%, P= 1.77E-07) leaflets both exhibited asymmetry. The C leaflet could also be distinguished from its mirror image, C’ (Fig. 5G&H, Accuracy=60.8%, P=0.028, Supplement Fig. 4F-J). This indicates that, unlike *E. crista-galli*, *C. speciosa* compound leaves are not bilaterally symmetric.

We also examined the leaves of *Chukrasia tabularis*, a large pinnate compound leaf, and found no mirror-symmetric relationship between leaflets on the left and right sides (Fig. 5I, P= 3.05E-09, Supplementary Fig. 5A-C), indicating *C. tabularis* compound leaves are also not bilaterally symmetric. We used similar methods to examine the pinnate compound leaves of *Dracontomelon duperreanum* and *Senna siamea*. Interestingly, *S. siamea*, like *C. tabularis*, exhibited bilateral asymmetry (Fig. 5J, P= 1.03E-08, Supplementary Fig. 5D-F), whereas *D. duperreanum* compound leaves were bilaterally symmetric (Supplementary Fig. 5G-I). Subsequently, we examined the leaves of some simple-leaf plants: *Ficus microcarpa*, *Triadica sebifera*, and *Talipariti tiliaceum*, finding their leaves all exhibited perfect bilateral symmetry (Fig. 5K-L, Supplementary Fig. 5J-L).

Fig. 5M shows the accuracy and statistical significance (-log10(P)) for distinguishing left and right halves of the leaves/sides mentioned above. This reveals that leaves can be either bilaterally symmetric or not; prior to rigorous verification, we cannot default to either assumption.

### Distinguishing Leaves from Stems of Different Chirality in *Mikania micrantha*

The chirality (handedness) of *Mikania micrantha* twining stems is variable; the same plant can simultaneously have left-handed (sinistrorse) and right-handed (dextrorse) twining stems, and chirality can switch [16, 17]. The mechanism determining chirality in *M. micrantha* remains unclear, but even if the determination involves random oscillation, it would entail oscillations in gene expression and potentially epigenetic modifications. We therefore hypothesized that such gene expression changes would also be recorded in leaf appearance. We collected leaves from dozens of *M. micrantha* plants from stems of both chiralities (L and R) for model training. Results also showed systematic differences between them (Fig. 6A-F, Supplementary Fig. 6), with identification accuracy exceeding 80% (P= 2.27E-07). The average images revealed subtle, visible differences between leaves from left-handed and right-handed stems (Fig. 6G), as expected. We then mirrored the R leaf images (R’) (Fig. 6G) and compared them with L leaves, finding they could still be successfully distinguished (Fig. 6H-J, Supplementary Fig. 6). Moreover, L vs R’ showed higher Accuracy, larger ROC AUC, and a smaller P-value (1.45E-08) compared to L vs R. This confirms our hypothesis: leaves from left-handed and right-handed twining stems are not only different but also more asymmetric. This result indicates that left-handed and right-handed twining stems are chiral opposites but do not possess detailed symmetry. This, too, was unverifiable before the introduction of AI.

**Fig. 6.**
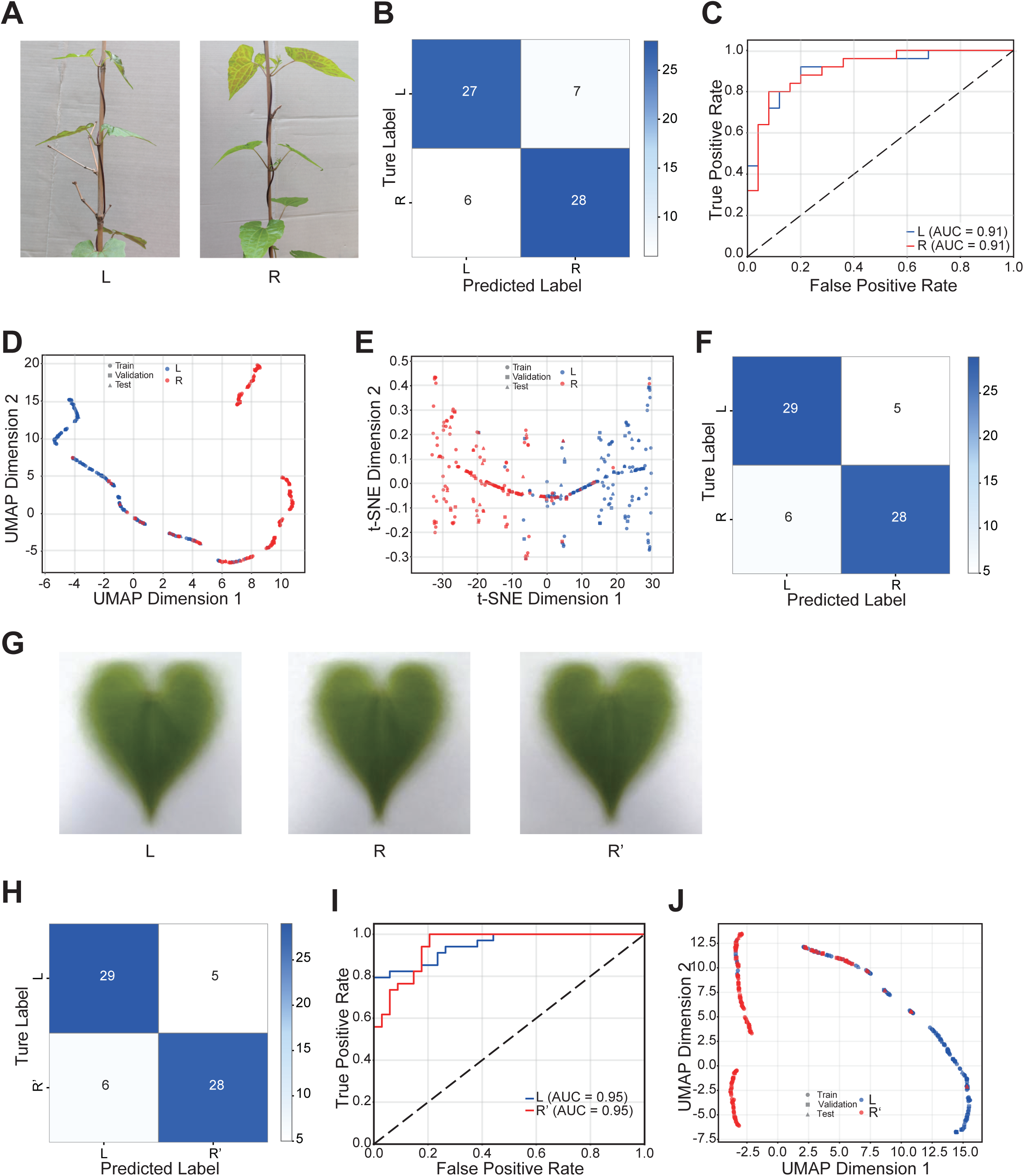
Comparison of leaf appearance from stems of different chirality in *Mikania micrantha*. A, Left-and right-handed twining stems of *M. micrantha*. B, Validation Accuracy for distinguishing leaves from left-vs. right-handed stems. C, ROC curve for the model classifying leaves from left-vs. right-handed stems. D, UMAP dimensionality reduction visualization of image similarity distributions. E, t-SNE dimensionality reduction visualization of image similarity distributions. F, Test Accuracy for distinguishing leaves from left vs. right-twining stems. G, Average images of leaves from left-handed stems (L), right-handed stems (R), and mirrored images of leaves from right-handed stems (R’). H, Validation Accuracy for distinguishing leaves from left-handed stems (L) vs. mirrored images of leaves from right-handed stems (R’). I, ROC curve for the model classifying leaves from L vs. R’. J, UMAP dimensionality reduction visualization of the image similarity distributions mentioned previously.

## Discussion

For humans, distinguishing facial features between individuals is crucial information [18]. Distinguishing organ appearance between different plant species is also relatively important (e.g., for identifying food vs. poison). However, distinguishing between individuals of the same species and cultivar seems unimportant, especially since plants are sessile – identifying them by location is far more economical. Consequently, humans did not evolve the ability to distinguish between two trees based on leaf appearance. For computers, however, there is no such distinction; humans, animals, and plants are all objective entities. Thus, a computer can perceive each plant’s leaves as unique, just as it perceives each human face as unique. Of course, before the advent of advanced computers and algorithms, we could not test this hypothesis. Fortunately, such work is now readily feasible.

This study not only validates our hypothesis but also offers several insights. First, it demonstrates that the characteristics of a plant’s genome and gene expression are indeed recorded in minute detail in leaf appearance, even though we cannot yet establish direct links between them – similar to our inability to strongly link a person’s facial features to their genome and physiological state. This “decoding” of organismal appearance is an important research direction poised for significant future progress. Second, “plant faces” hold potential application value in ecology, agronomy, plant protection, and plant development, such as tracking individual plant growth, mapping invasion routes of invasive plants, and creating facial profile archives for trees to protect rare species or ancient trees.

The identification of compound leaflet positions and leaves from stems of different chirality share similarities with but also differ from the typical “plant face” concept. Compound leaf development has long been a key topic in developmental botany [19]. Distinguishing leaflets links their appearance to concentration gradients of relevant substances during development [20], potentially providing insights and assistance for future research on compound leaf development. People might not have anticipated systematic appearance differences between leaves from different individual plants of the same cultivar, but they might readily imagine systematic differences among leaflets at different positions on a compound leaf. However, experiments prove that the former differences are far more significant than the latter. This misperception arises because when observing compound leaflets, people usually reference their positions, and leaflets at different positions often differ in size and shape. But if these leaflets are photographed independently, human eyes also struggle to distinguish them (e.g., a smaller leaflet from a large compound leaf might be larger than a larger leaflet from a small compound leaf). The disparity between computer and human recognition also highlights the unreliability of human intuition, necessitating experimental verification.

In common perception, aside from a few exceptions, most plant leaves are statistically expected to be bilaterally symmetric, although individual leaves may not be perfectly symmetric. Without using computers, this remains an assumption, lacking evidence. Through machine learning validation, we found that leaves can indeed be bilaterally symmetric, but they may also not be. Before verification, it is difficult to predict whether a plant’s leaves are symmetric. Our preliminary results suggest that simple-shaped simple leaves tend to be symmetric, whereas compound leaves, especially complex ones, are more prone to asymmetry. This implies that the more complex the leaf structure, the more complex its developmental regulation, and the harder it is to achieve perfect symmetry.

Stem chirality is another long-standing fascinating topic, discussed since Darwin’s time [21]. However, research on the determination of twining chirality remains scarce, primarily due to a lack of effective methods. Theoretically, gene expression must differ between the two chiralities of Variable Chirality Twining Stems (VCTS) [16], yet their appearance seems perfectly symmetric to the human eye, creating a sense of contradiction. Now, through computer recognition, we know that leaves from stems of different chirality indeed exhibit subtle differences, confirming the hypothesis of differential gene expression. Comparing the transcriptomes and epigenetic modifications of stems with different chirality might be a feasible approach to explore the mechanism behind twining chirality determination.

The differences in leaves from stems of different chirality share some similarity with compound leaf asymmetry. They suggest a possibility: due to the chiral nature of fundamental biological building blocks – sugars, amino acids, phospholipids, proteins, and nucleic acids are all chiral molecules – achieving perfect bilateral symmetry in development is challenging. Perhaps organisms can only achieve approximate bilateral symmetry, and when observed with high-resolution tools like AI, many asymmetric details become apparent.

It is worth emphasizing that our research did not require specialized photographic equipment or large computers, nor GPUs; it was conducted using ordinary smartphones and personal computers. This reduces economic and time costs, facilitating larger-scale studies within the same timeframe. Even ***amateurs*** can readily begin experiments if interested. Naturally, using professional equipment would undoubtedly yield better results. The preliminary success of the “plant face recognition machine” suggests the potential for developing networked applications, like smartphone APPs, enabling dynamic enrollment and detection via smartphones and other devices, greatly expanding future application prospects and enhancing the application potential of leaf appearance as a vital component of the phenome.

In conclusion, this study not only demonstrates the “Plant Face” concept but also highlights the immense value of deep learning as a powerful “sensory extension” tool for discovering novel biological principles. Our work provides new research ideas and technical pathways for plant individual biology, developmental biology, and invasive plant ecology (using *M. micrantha* as a model). The current research is a preliminary attempt, and we hope more colleagues will join in, studying more species, validating the plant face hypothesis, and expanding its applications.

## Materials and Methods

### Plant Materials

The plant specimens used in this study (see Supplementary Table) were collected primarily from Haizhu District and Huangpu District in Guangzhou, Guangdong Province, China, with the exception of peach (*Prunus persica*) and some paper mulberry (*Broussonetia papyrifera*) leaves, which were collected from North China. All collections were conducted between June and October. The selected plants included common horticultural varieties, invasive species, and widespread wild plants. Importantly, no rare or protected wild plant species were involved in this study.

### Equipment

Digital photographs were captured using various commercially available smartphone models, including Huawei, Xiaomi, and iPhone devices, all purchased from the Chinese consumer market. The computational programs, written in Python, did not require specialized hardware and could execute efficiently on standard personal computers or servers without GPU acceleration.

### Collection and Imaging Protocol

Leaves were carefully excised from plants, placed in plastic bags, and transported to the laboratory for imaging. A standardized imaging setup was employed: white paper served as the background, with leaves oriented vertically. When leaves could not lie flat naturally, white modeling clay was used as an adjustable support. Photographs were taken by hand-holding smartphones, with the distance adjusted to maximize the leaf area within the frame while ensuring complete capture of the leaf blade (excluding the petiole). The macro mode was deliberately disabled on all devices to maintain consistent imaging conditions across samples.

### Image Preprocessing

Minimal preprocessing was applied to the captured images. For photographs with excessive marginal blanks, the empty borders were cropped while strictly maintaining the original aspect ratio of the images.

## Mathematical Foundations

### Convolutional Neural Networks

The core architecture employs convolutional operations defined as:

### Discrete Convolution

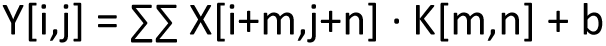

where X is the input feature map, K is the convolution kernel, and b is the bias term.

### Activation Function (ReLU)

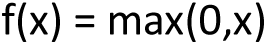

Rectified Linear Units introduce non-linearity while maintaining gradient propagation efficiency.

### MobileNetV2 Architecture **[22]**

The system utilizes depthwise separable convolution to reduce computational complexity:

### Standard Convolution Complexity

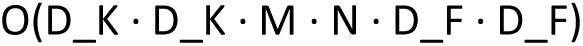

### Depthwise Separable Convolution Complexity

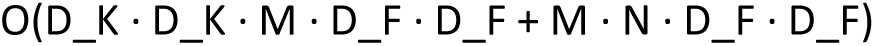

where D_K is kernel size, M is input channels, N is output channels, and D_F is feature map size.

### Inverted Residual Blocks

MobileNetV2 employs bottleneck layers with expansion factor t:

- Expansion: M → t·M channels via 1×1 convolution
- Depthwise convolution: t·M → t·M channels
- Projection: t·M → N channels via 1×1 convolution

### Similarity Metrics and Loss Functions Cosine Similarity

For feature matching in the recognition system:

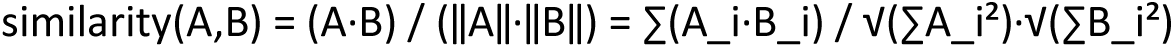

### Categorical Cross-Entropy Loss

For multi-class classification:

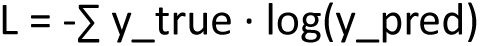

where y_true is the one-hot encoded true label and y_pred is the predicted probability distribution.

## Dimensionality Reduction Techniques

### t-SNE (t-Distributed Stochastic Neighbor Embedding) [23]

Minimizes Kullback-Leibler divergence between high-dimensional and low-dimensional distributions:

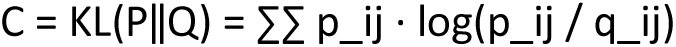

### UMAP (Uniform Manifold Approximation and Projection) [24]

Optimizes fuzzy topological structure preservation using cross-entropy:

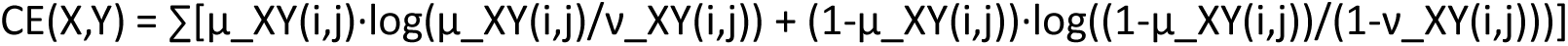

### Programming and System Development

Codes are available at git-hub.

### Image Classification System

(https://gist.github.com/MarchLiu/f51f472969ab37ec2b22d1e38fb57f75)

The automated leaf classification system was developed using Python 3.8 with TensorFlow 2.5.0 as the primary deep learning framework. Key programming aspects included:

### Data Pipeline Development

- Implemented an automated data splitting class (DataSplitter) that partitions image datasets into training (60%), validation (20%), and test (20%) subsets while maintaining class balance
- Created custom data generators with real-time augmentation including rotation (±8°), translation (±13%), zoom (±10%), and brightness adjustment (90-110%)
- Established separate analysis generators with fixed random seeds to ensure reproducible feature extraction

### Model Architecture Implementation

- Built a transfer learning framework using MobileNetV2 as the feature extractor with frozen weights
- Added custom classification head with dense layers (1024 → 512 → 256 → 128 units) and dropout regularization (30-50%)
- Implemented early stopping with 10-epoch patience and model checkpointing to prevent overfitting

### Debugging and Optimization

- Addressed TensorFlow compatibility issues by setting specific environment variables and random seeds
- Implemented comprehensive error handling for missing dependencies and corrupted image files
- Developed visualization utilities for training curves, confusion matrices, and feature space projections
- Optimized batch processing and memory usage to handle high-resolution leaf images (224×224×3)

### Plant Recognition System

(https://gist.github.com/MarchLiu/b1a4e1504a5b1d4ee97a3a92f56166a3)

The plant face recognition system employed a different architectural approach focused on individual plant identification:

### Database Management

- Designed a JSON-based plant database with feature vector storage using Base64 encoding
- Implemented duplicate detection through MD5 file hashing to prevent data redundancy
- Created statistical tracking for recognition accuracy across different database sizes

### Feature Extraction Pipeline

- Developed robust model persistence with automatic fallback mechanisms when primary loading failed
- Implemented cosine similarity-based matching with configurable confidence thresholds (default: 0.7)
- Added feature vector normalization (L2 normalization) to improve distance metric reliability

### System Integration

- Created interactive command-line interface with multiple operational modes (enrollment, recognition, validation)
- Implemented administrator confirmation workflow with optional feature database updates
- Developed comprehensive statistical analysis for system performance evaluation

### Implementation Details

Both systems were developed with emphasis on reproducibility through fixed random seeds, comprehensive logging, and version-controlled dependencies. The modular architecture allows for easy extension and adaptation to different plant species and imaging conditions. All mathematical operations were implemented using NumPy and TensorFlow optimized functions to ensure computational efficiency and numerical stability.

The debugging process involved extensive validation of feature extraction consistency, database integrity checks, and performance monitoring across different hardware configurations to ensure robust operation in research environments.

## Supporting information

Main text with Figures

## Author Contributions

Y.G. conceived the original idea. Y.G., Y.Z., X.L., and Y.J.Z. discussed and designed the research. J.L., Y.C., Z.G., X.Z., Z.X., G.M., and Y.G. collected leaf samples and performed photography. Y.G., X.L., J.L., and Y.C. developed and debugged the computational programs. X.L. and Y.C. were responsible for the maintenance of computational software and hardware. All authors participated in discussing and summarizing the experimental results. Y.G., J.L., X.L., and Y.Z. wrote the manuscript draft. Y.G. supervised the study.

## Acknowledgement

We are grateful to Mr. Gehan Huang from Suzhou Customs for his valuable support in the identification of species.

