## Supplementary material for "Plant Face: Machine learning decodes genetic, environmental and developmental imprints in leaf appearance": Main text with Figures

**Supplementary Table 1.**

The plants involved in this study.

| species | number of plants |
|---|---|
| *Erythrina crista-galli* | 14 |
| *Triadica sebifera* | 6 |
| *Osmanthus fragrans* | 4 |
| *Broussonetia papyrifera* | 3 |
| *Ficus microcarpa* | 3 |
| *Loropetalum chinense var. rubrum* | 2 |
| *Fagraea ceilanica* | 2 |
| *Prunus persica* | 2 |
| *Ficus virens* | 2 |
| *Chukrasia tabularis* | 2 |
| *Manilkara zapota* | 2 |
| *Talipariti tiliaceum* | 2 |
| *Ficus stenophylla* | 1 |
| *Antidesma bunius* | 1 |
| *Olea europaea* | 1 |
| *Dracontomelon duperreanum* | 1 |
| *Senna siamea* | 1 |
| *Ilex rotunda* | 1 |
| Sum. | 50 |
| *Ceiba speciosa* | >6* |
| *Mikania micrantha* | >30* |

* Not for face recognition, only for the studies on compound leaf and twining stem.

**Supplementary Figure Legends**

**Supplementary Fig. 1.**
More comparison of leaves from two plants of *Erythrina crista-galli*. A & B, The training history for the classification between #0 and #2 (A) and between #1 and #2 (B). C-E, the Validation confusion matrix (C), the ROC curve (D) and the Test confusion matrix (E) of the classification of #0 and #2. F-H, the Validation confusion matrix (F), the ROC curve (G) and the Test confusion matrix (H) of the classification of #1 and #2.

**Supplementary Fig. 2.** Training performance and evaluation of the ten-class model. A, Training curves depicting the evolution of model accuracy (A) and loss (B) throughout the learning process relative to epochs. C, Comparative class-specific accuracy between validation and test datasets. D, Class-wise performance metrics for 10 classes.

**Supplementary Fig. 3.** Training dynamics and position-specific evaluation in *E.crista-galli* and *C.speciosa* leaf classification. A-E, Training history and validation analysis for *E.crista-galli* model. A, Training history: changes in Accuracy (A) and Loss (B) relative to epochs. C, UMAP visualization of image feature similarity distributions. D, Position-wise classification metrics for 3 positions. E, Validation vs. Test accuracy comparison per position. F-J, Training performance and validation analysis for *C.speciosa* model. F, Training history: changes in Accuracy (F) and Loss (G) relative to epochs. H, UMAP projection of image feature similarities. I, Position-specific validation vs. test accuracy. J, Performance metrics across 5 positions.

**Supplementary Fig. 4.** t-SNE visualization and ROC analysis of leaflet laterality classification in *E. crista-galli* and *C.speciosa*. A, t-SNE projection illustrating latent feature similarity distributions between left (L) and right (R) leaflets. B, ROC curve for L vs. R' discrimination in *E.crista-galli* (R' = mirrored R). C, ROC curve of innate vs. mirrored medial leaflet discriminability (*E.crista-galli* C, innate; C', mirrored). D, t-SNE projection of image features differentiating lateral (D: L vs. R') and medial (E: C vs. C') in *E. crista-galli*. F, ROC curve for distinguishing *C.speciosa* L2 vs. R2' (the mirrored R2) (F) and L1 vs. R1 (the mirrored R1) (G). H, t-SNE visualization of *C.speciosa* leaflet feature distributions: L2 vs. R2' (the mirrored R2) (H), L1 vs. R1' (the mirrored R1) (I), and C vs. C' ( the mirrored C) (J).

**Supplementary Fig. 5**. Bilateral symmetry discriminability in compound and simple leaves: ROC/t-SNE analysis of innate versus mirrored leaflets across six species. A, D, G. Representative images of pinnate compound leaves of *Chukrasia tabularis* (A), *Senna siamea* (D), and *Dracontomelon duperreanum* (G). (L represents the left compound leaf, R represents the right compound leaf).

B, E, H. ROC curves assessing discriminability between native left leaflets (L) and mirror-flipped right leaflets (R') in: *C.tabularis* (B), *S.siamea* (E), and *D.duperreanum* (H). C, F, I. t-SNE visualization of lateral leaflet symmetry distributions (L: native left vs. R': mirrored right) in: *C.tabularis* (C), *S.siamea* (F), and *D.duperreanum* (I). J-L, ROC curves evaluating classifier performance in disscriminating original (0) vs. mirror-flipped (0') leaf images for simple-leaf species: *F.microcarpa* (J), *T.sebifera* (K), and *T.tiliaceum* (L).

**Supplementary Fig. 6.** Machine learning performance in classifying leaf laterality (L/R) and mirrored-flipped leaves (L/R') in *Mikania micrantha*. A, Training dynamics for L vs. R classification: Accuracy (A) and Loss (B) across epochs. C, Validation vs. test accuracy per class (L vs. R). D, Class-wise performance metrics (L vs. R). E, Training dynamics for L vs. R' classification : Accuracy (E) and Loss (F) across epochs. G, Confusion matrix for test data (L vs. R'). H, Class-wise performance metrics (L vs. R'). I, Validation vs. test accuracy per class (L vs. R').

### Supplementary Fig. 1

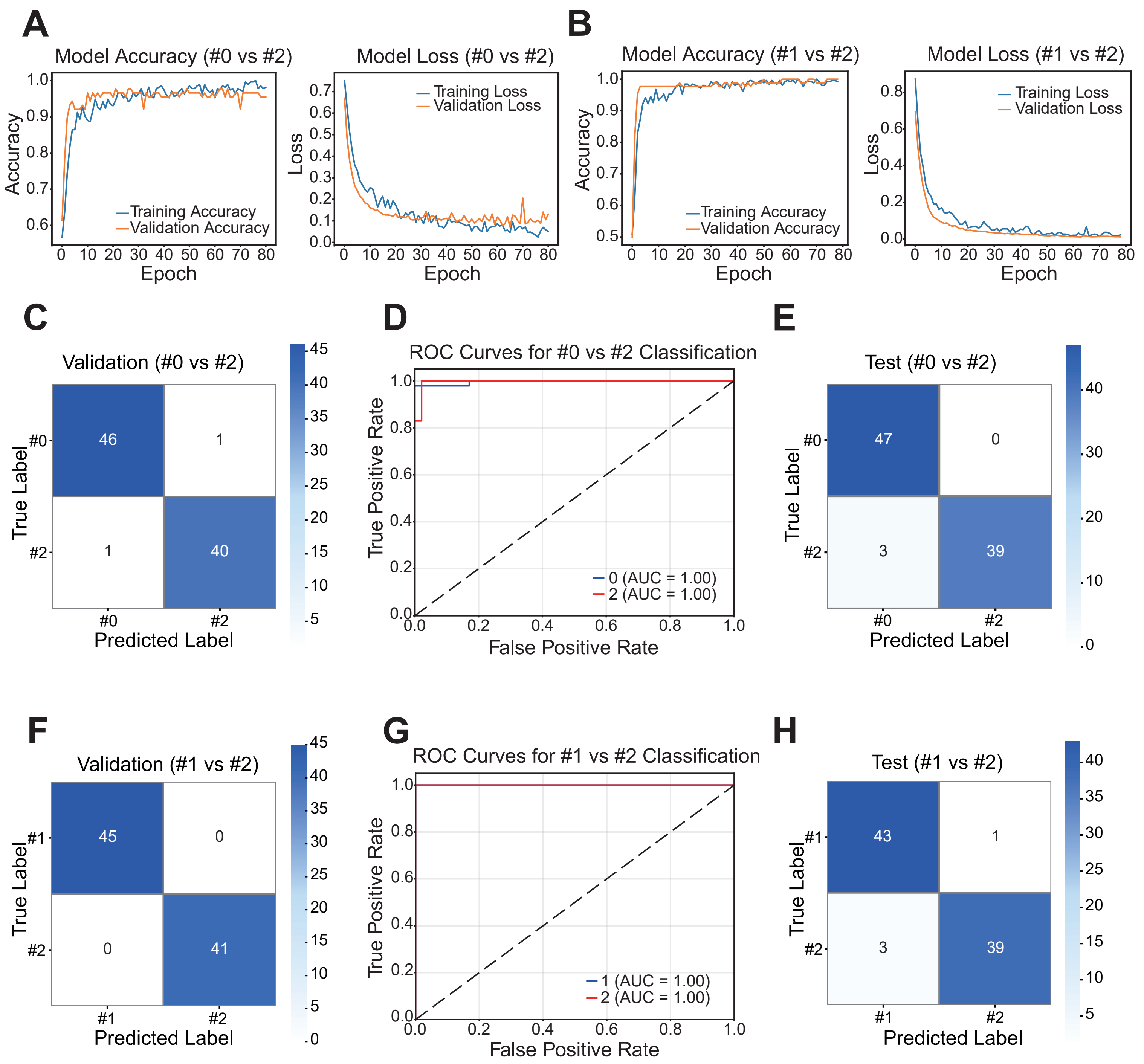

### Supplementary Fig. 2

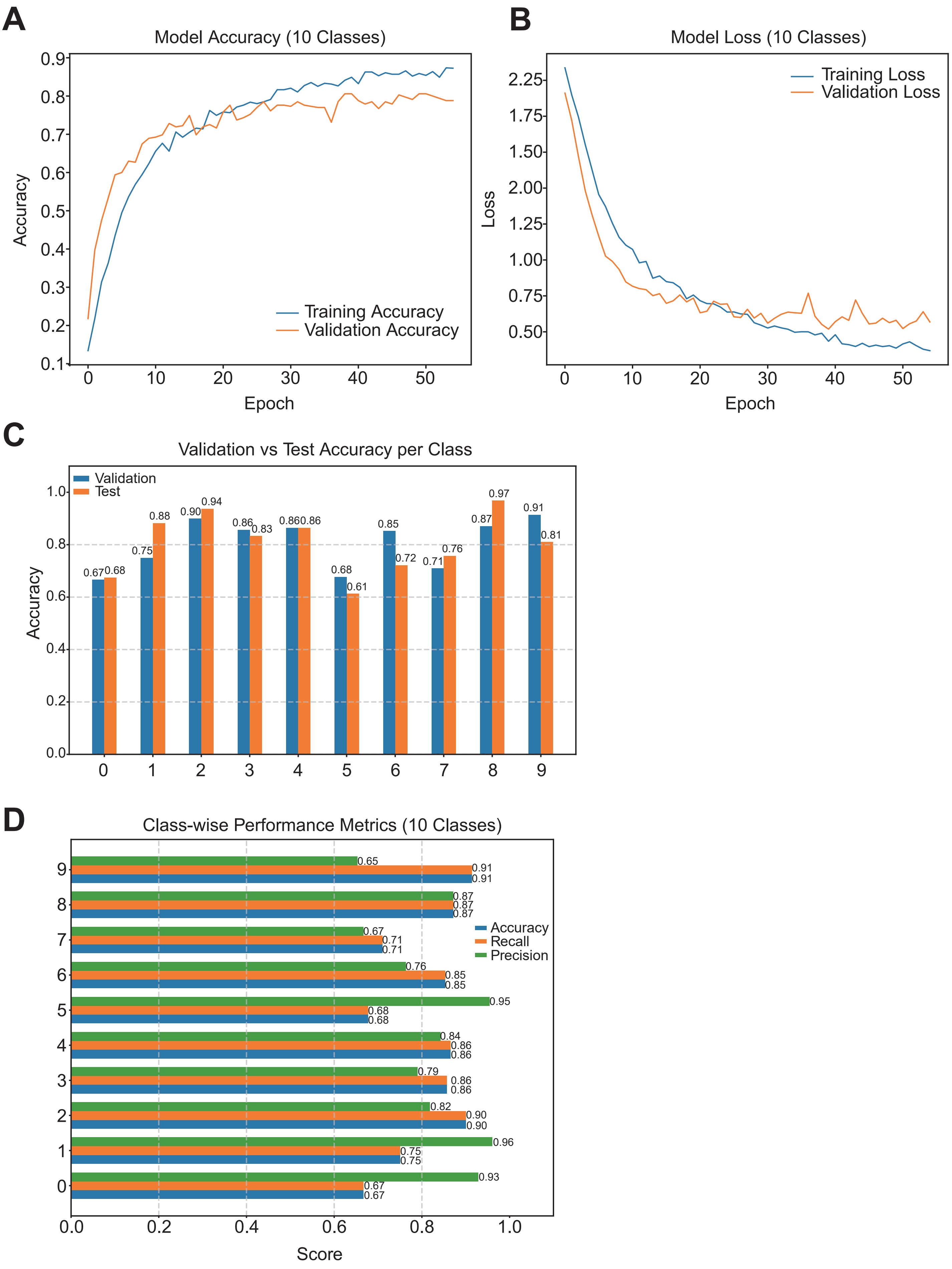

### Supplementary Fig. 3

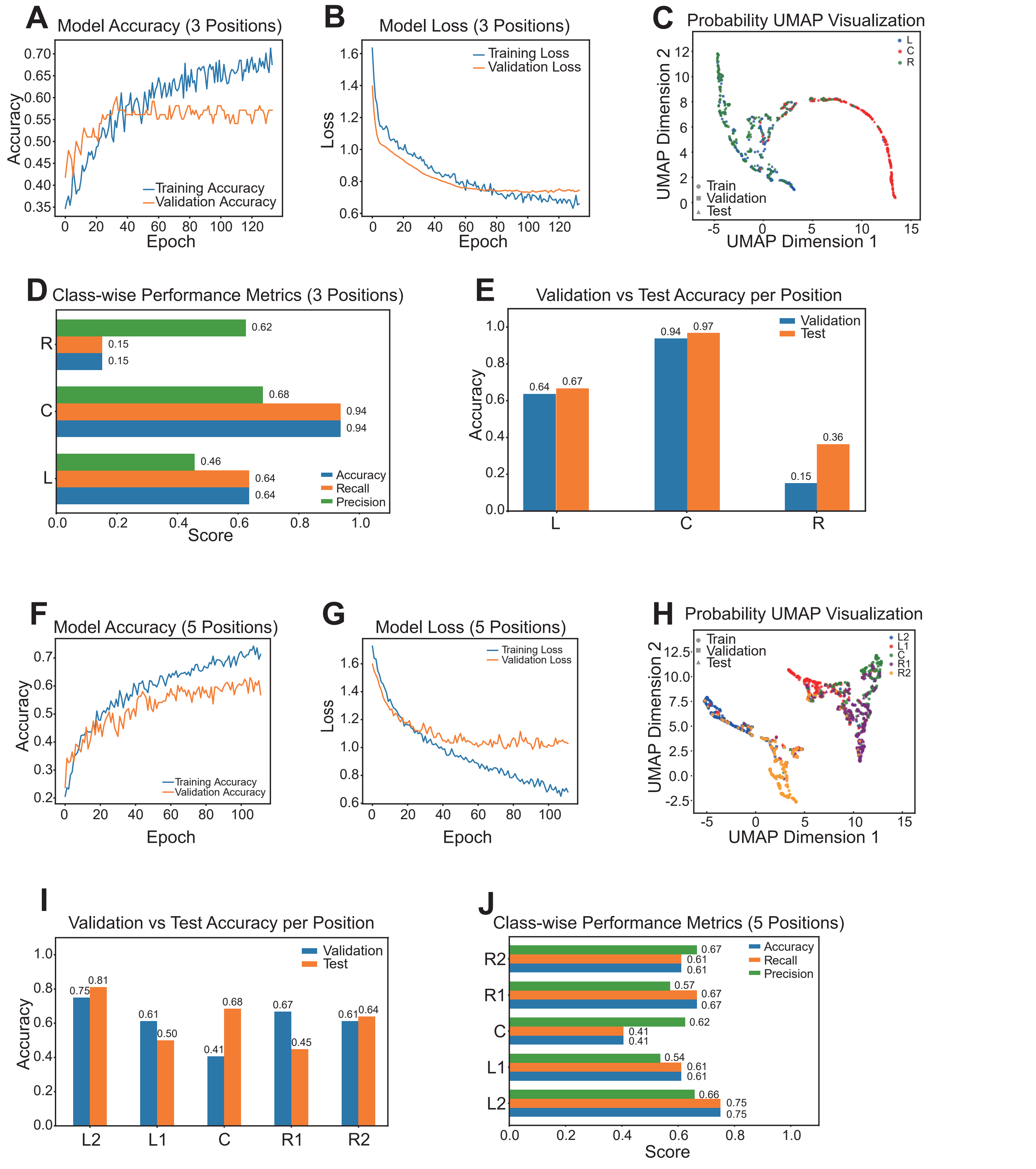

### Supplementary Fig. 4

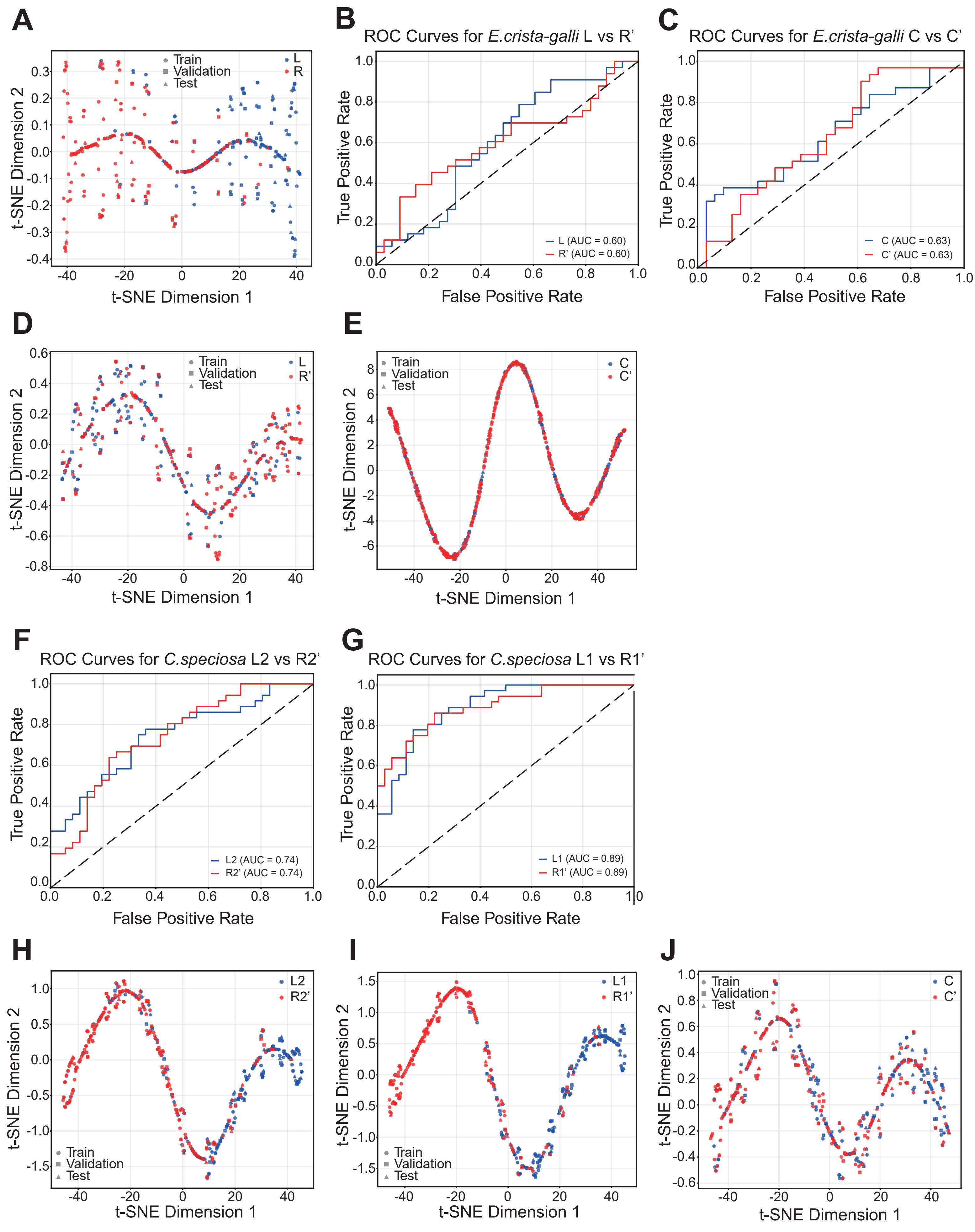

### Supplementary Fig. 5

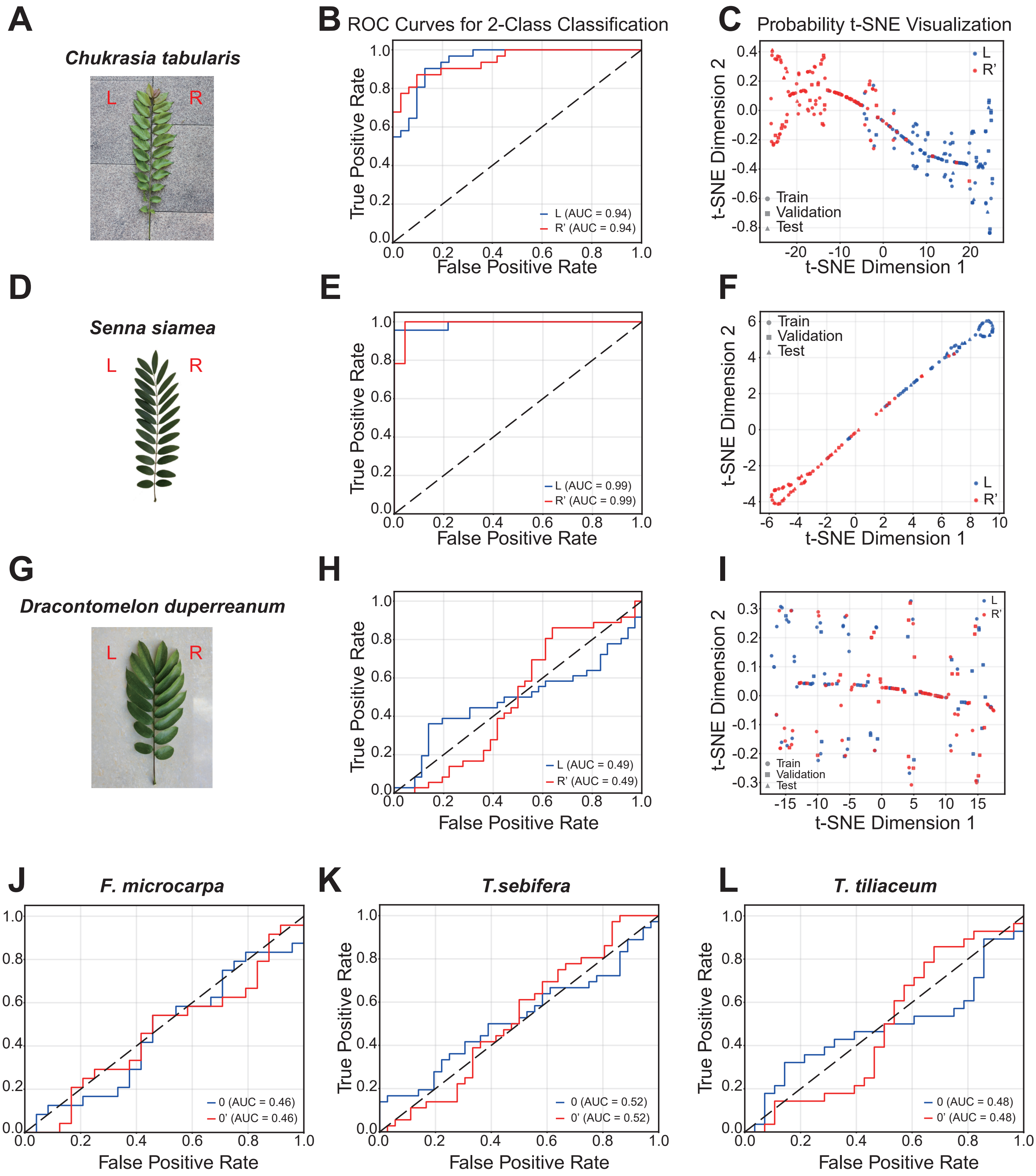

### Supplementary Fig. 6

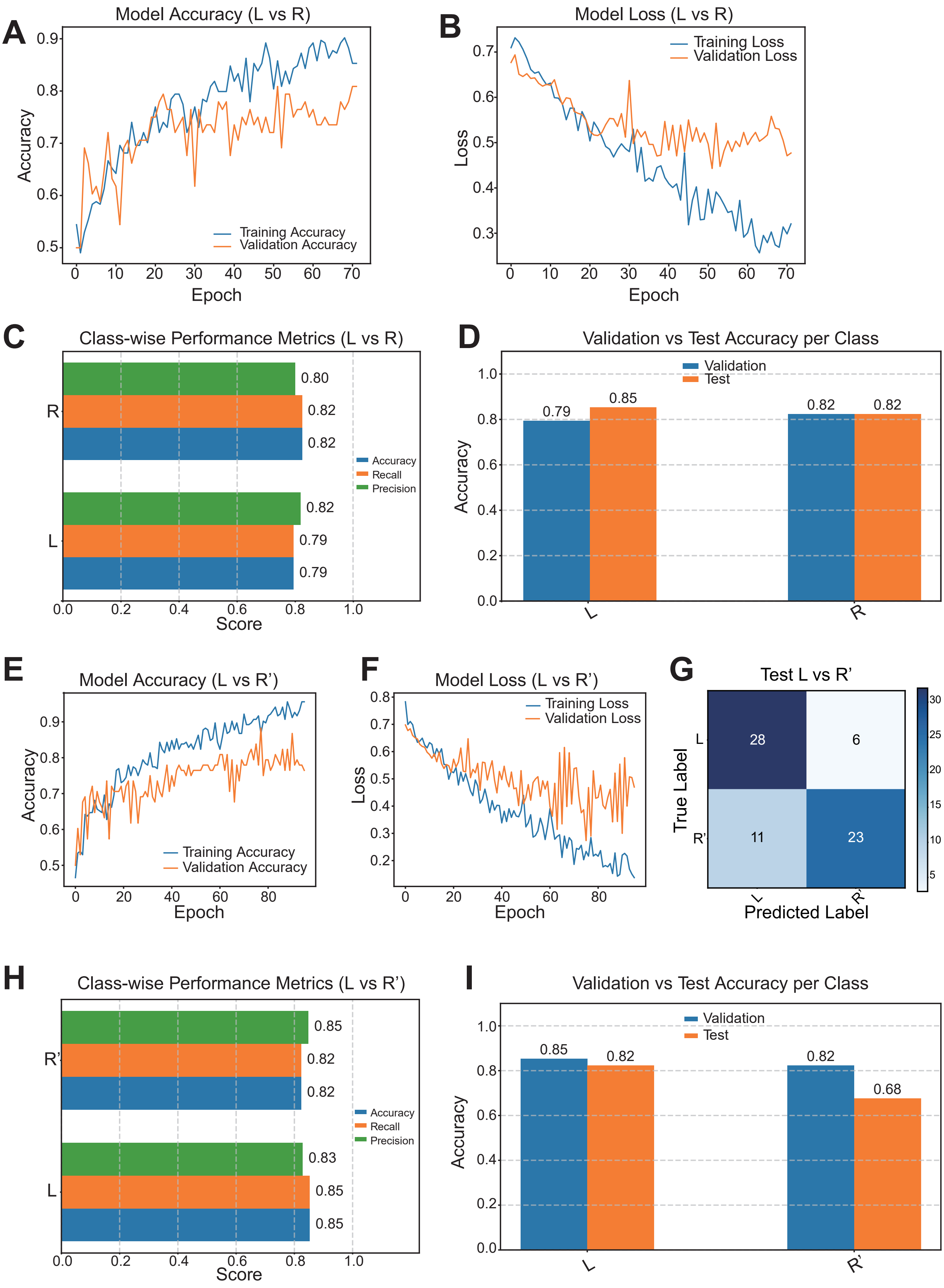
